# Molecular convergence and positive selection associated with the evolution of symbiont transmission mode in stony corals

**DOI:** 10.1101/489492

**Authors:** Groves B. Dixon, Carly D. Kenkel

## Abstract

Heritable symbioses are thought to be important for the maintenance of mutually beneficial relationships (1), and for facilitating major transitions in individuality, such as the evolution of the eukaryotic cell (2, 3). In stony corals, vertical transmission has evolved repeatedly (4), providing a unique opportunity to investigate the genomic basis of this complex trait. We conducted a comparative analysis of 25 coral transcriptomes to identify orthologous genes exhibiting both signatures of positive selection and convergent amino acid substitutions in vertically transmitting lineages. The frequency of convergence events tends to be higher among vertically transmitting lineages, consistent with the proposed role of natural selection in driving the evolution of convergent transmission mode phenotypes (5). Of the 10,774 total orthologous genes identified, 403 exhibited at least one molecular convergence event and evidence of positive selection in at least one vertically transmitting lineage. Functional enrichments among these top candidate genes include processes previously implicated in mediating the coral-*Symbiodiniaceae* symbiosis including endocytosis, immune response, cytoskeletal protein binding and cytoplasmic membrane-bounded vesicles (6). We also identified 100 genes showing evidence of positive selection at the particular convergence event. Among these, we identified several novel candidate genes, highlighting the value of our approach for generating new insight into the mechanistic basis of the coral symbiosis, in addition to uncovering host mechanisms associated with the evolution of heritable symbioses.

**DATA ARCHIVAL LOCATION:** Raw sequencing data generated for this study have been uploaded to NCBI’s SRA: PRJNA395352. All bioinformatic scripts and input files can be found at https://github.com/grovesdixon/convergent_evo_coral.

## Introduction

For organisms that engage in symbiosis, the mode in which symbionts are transmitted to the next host generation is a major factor governing the ecological and evolutionary dynamics of the relationship across multiple scales of biological organization. For example, transmission mode is known to influence genome size and content, cooperative interactions between partners, holobiont ecology, and the speciation rates of both partners (7–11). Two transmission modes predominate in nature: offspring can either directly inherit symbionts, typically through the maternal line in the process of vertical transmission, or they can horizontally acquire symbionts from the environment, usually early in their development (reviewed in (12)); although it is important to note that the mode of transmission can change over evolutionary time (13) and mixed-mode transmission is also possible (12). In microbial symbioses, horizontal transmission is the basal state and repeated transitions to vertical transmission may have arisen as a means to further promote host-symbiont cooperation (13–15). Vertical transmission has been hypothesized to play an important role in the maintenance of mutually beneficial symbioses (1), and likely facilitated major evolutionary transitions in individuality, such as the evolution of the eukaryotic cell (2, 3). From the perspective of the symbiont, the genomic consequences of evolving a heritable symbiosis include a reduction in genome size and increased dependence on their hosts due to the loss of functionally redundant genes (3, 10). However, the underlying genetic architecture facilitating evolution of a heritable symbiosis from the perspective of the host remains unresolved.

The evolution of vertical transmission is predicted to be correlated with the evolution of host control mechanisms (16) and theory predicts a high rate of mutation in genes responsible for the host-symbiont fitness interaction (17). Selection on mechanisms critical for the establishment and maintenance of a horizontally transmitted symbiosis, such as cell surface molecules mediating inter-partner recognition, is likely also relaxed (12). Among metazoan hosts, diverse behavioral, developmental and physiological mechanisms are known to facilitate the vertical transmission of microbial endosymbionts (13, 16), yet there is also some evidence for phenotypic convergence. For example, plant-sucking stinkbugs and lice require microbial gut symbionts to facilitate digestion of sap and blood, respectively, but both have evolved additional specialized organs for housing bacteria along the female reproductive tract for the transmission of symbionts to eggs (16, 18). Convergent evolution at the phenotypic level is often the result of similar changes at the genomic level (19, 20) and comparative analyses have facilitated understanding of the genetic basis of convergently evolved phenotypes in diverse taxa (19, 21, 22). Therefore, by comparing vertically transmitting lineages with their closest horizontally transmitting relatives it may be possible to identify candidate genes involved in the evolution of convergent transmission mode phenotypes.

Reef-building corals exhibit both horizontal and vertical transmission of their obligate intracellular *Symbiodiniaceae* symbionts, offering an ideal opportunity to utilize such a comparative approach to identify candidate genes involved the evolution of symbiont transmission mode. The majority of coral species acquire their symbionts from the environment early in their development, but vertical transmission is exhibited by species in multiple different lineages, indicating that transmission mode has evolved independently at least four times (4, 23). Yet there is also significant morphological, physiological and ecological trait variation across the coral phylogeny (24), which can confound a comparative approach. In corals, transmission mode is often correlated with reproductive mode as coral species which broadcast spawn gametes tend to exhibit horizontal transmission, while species that internally brood larvae largely transmit symbionts vertically (4). However, the association is not perfect; some *Porites* spp. and all known *Montipora* spp. have convergently evolved to broadcast spawn eggs which contain *Symbiodiniaceae* (25, 26). We therefore sequenced the transcriptome of the vertically transmitting broadcast spawner, *Montipora aequituberculata*, in addition to mining other publicly available coral sequence resources (Table S1), to compile a set of transcriptomic references in which vertical transmitters could be compared with their closest horizontally transmitting relatives, while also accounting for variation in other life-history traits (Fig. 1). From this dataset, we inferred orthologous groups and identified genes showing both signatures of positive selection and convergent amino acid substitutions (overlapping amino acid changes resulting from independent amino acid substitutions at the same position in two or more lineages). We found that the frequency of molecular convergence tended to be higher among vertically transmitting lineages and although top genes are enriched for biological processes previously implicated in the coral-algal symbiosis, we also identify several novel candidates, generating new insight into the mechanistic basis of this relationship.

**Figure 1.**
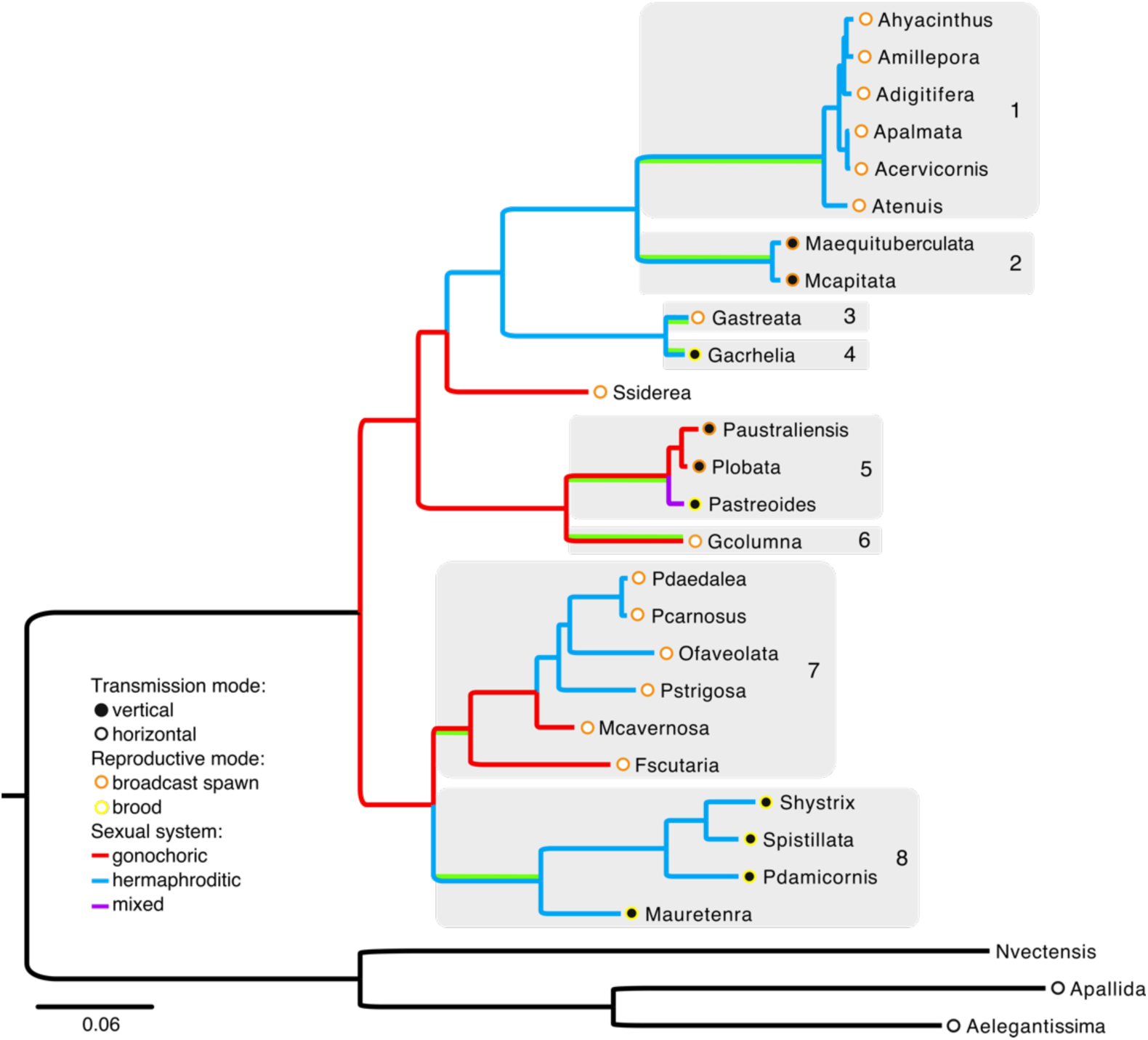
Species tree with phenotypic labels indicating transmission mode, reproductive mode and sexual system (4, 26, 94). Vertically transmitting species are indicated by filled circles at their terminal nodes, horizontally transmitting species with open circles at their terminal nodes. For each clade (1-8), the particular branch examined for convergent substitutions and positive selection is indicated by a green highlight. In each case, this is the branch leading the common ancestor of the clade. Shaded clades were considered when describing overlapping convergence events, referred to as (1) Sister *Montipora*, (2) *Montipora*, (3) Sister *Galaxia*, (4) *Galaxia*, (5) *Porites*, (6) Sister *Porites*, (7) Sister Pocilloporid, (8) Pocilloporid.

## Results & Discussion

### Ortholog identification

To examine molecular convergence and positive selection, we compared homologous coding sequences from transcriptomic data of 25 coral species. First, protein coding sequences were predicted from the transcriptomic data based on open reading frames and sequence homology to known proteins (27) and protein domains (28), and *FastOrtho* (29) was used to assign sequences to preliminary orthologous groups (N = 106,300 groups). A subset of 1,196 single-copy orthologous groups with at least 20 of the 28 taxa represented was used to construct a species tree (Fig. 1), which recapitulates known relationships reported in earlier studies using single-gene (23, 30) and multi-gene phylogenies (31). We then identified putative single-copy orthologs (groups with only a single representative sequence from each species) from the initial set of 20,563 orthologous groups for which at least 7 (25%) of the species were represented. Of these, 9,794 were truly single copy, whereas 10,769 had multiple sequences for one or more species. Two biological explanations for this observation are gene duplication events subsequent to the relevant speciation event, or transcript isoforms of the same gene (32). Transcript isoforms are more likely given the nature of the dataset, but in either case, any sequence from these monophyletic groupings can be appropriately compared to those from other species. Therefore, rather than eliminate all orthologous groups with multiple sequences, we applied a filtering approach similar to that described by (32) to retain an additional 3,298 of the 10,769 multiple sequence orthologs. Specifically, we constructed gene trees from the protein alignments and pruned away all but the longest of multiple sequences from single species that formed monophyletic clades (Fig. S1; see Methods). In this way, we identified a total of 13,092 total single copy orthologs. Orthologs were then aligned using MAFFT (33) and reverse translated into codon sequences using Pal2Nal (34).

Orthologs were further quality filtered based on monophyly of known clades. Individual gene trees were constructed from nucleotide alignments of each single-copy ortholog and checked for monophyly of known clades (Fig. 1, 1-8 and Robusta/Complexa). All species fell within their expected clades in 58% of the gene trees. If a single sequence fell outside of its expected clade or clades, that sequence was removed and the ortholog was retained (27% of orthologous groups). If more than one sequence fell outside its expected clade the ortholog was removed (15% of orthologous groups). In total, this left 119,049 sequences (mean species per orthologous group = 10.7) comprising 11,130 orthologous groups, hereafter referred to as genes, which were used for the ancestral reconstruction and branch-site tests. Genes with fewer than 5 representative sequences were also removed, resulting in a final total of 10,774 genes.

### Evidence of positive selection and molecular convergence

For each orthologous nucleotide alignment, PAML (35) was used to reconstruct the ancestral amino acid at each node in the species tree and identify the amino acid changes that occurred along the branches of the tree. We focused our analysis on eight clades (four with vertical transmission and four with horizontal transmission), and identified all overlapping substitutions, or independent substitutions occurring at the same position between the branches leading directly to these clades’ most recent common ancestors (Fig. 1). We classified substitutions according to the type of change observed: parallel substitutions refer to the same derived amino acid evolving from the same ancestral amino acid, convergent substitutions refer to same derived amino acid evolving from different ancestral amino acids, divergent substitutions refer to different derived amino acids evolving from the same ancestral amino acid and ‘all different’ refer to different derived amino acids evolving from different ancestral amino acids. Following (36), we consider both parallel and convergent substitutions to be indicative of molecular convergence.

Among the vertical transmitters, we identified 8,952 amino acid positions exhibiting either parallel (n=8,877) or convergent (n=75) substitutions in at least two lineages (ancestral reconstruction posterior estimate > 0.8, Fig. 2A, Fig. S2). The convergence events occurred in 4,117 out of 10,774 total genes in the dataset, with an average of 0.71 convergent sites identified per gene (median = 0; Fig. 2B). Of the four possible types of overlapping substitutions, convergent substitutions were by far the least frequent (Fig. 2A; Fig. S2). The most common type was divergent substitutions. The two remaining types, parallel and ‘all different’ occurred with roughly similar frequency (Fig. 2A). Across the entire dataset, 11% of overlapping substitutions were classified as molecular convergence (convergent or parallel).

**Figure 2.**
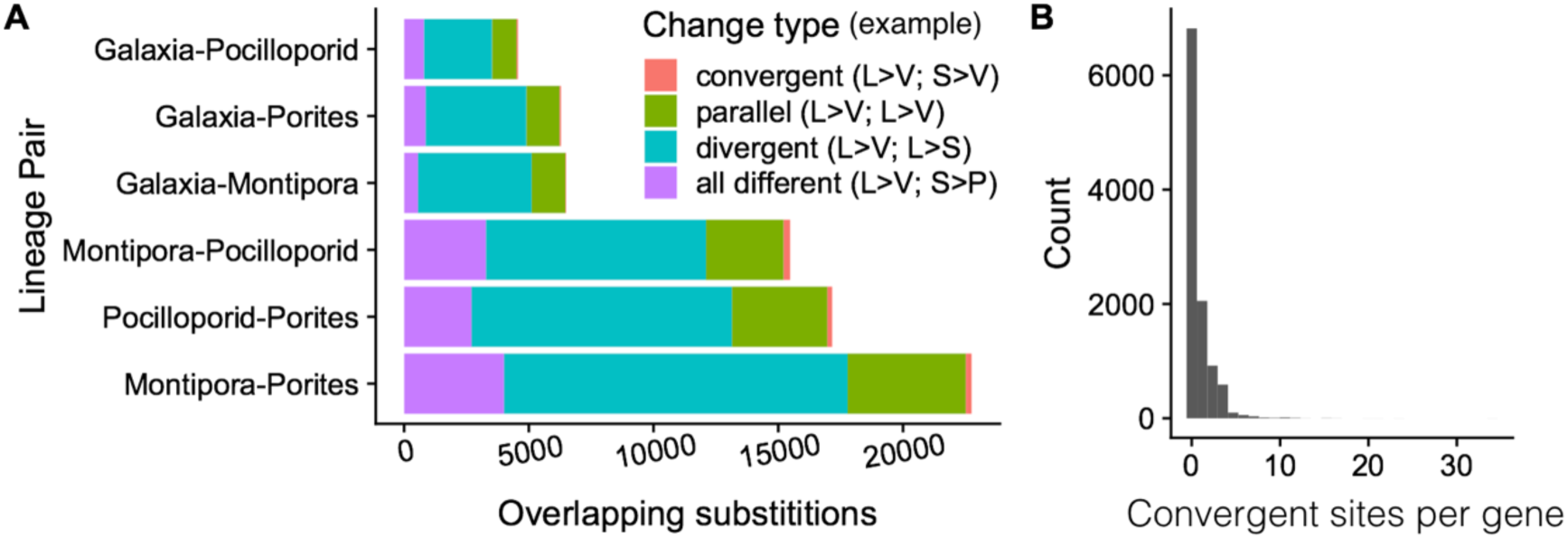
Frequency of convergence events. (A) An overlapping substitution is defined as an inferred amino acid change that occurred at the same position independently in the lineages leading to the common ancestor of the two indicated vertically transmitting clades. Each overlapping substitution was classified into one of four categories: convergent substitutions (least frequent; salmon) are changes from different amino acids to the same amino acid; parallel substitutions (second most frequent; green) are changes from the same amino acid to the same new amino acid; divergent substitutions (most common; teal) are changes from the same amino acid to a different one; ‘all different’ substitutions (third most common; purple) are changes from different amino acids to different new amino acids. (B) Histogram of the number of sites showing molecular convergence (convergent or parallel substitutions) per tested gene (mean = 0.71; median=0).

In addition to quantifying molecular convergence, we also tested for evidence of positive selection in each vertically transmitting lineage and for all vertically transmitting lineages at once using the branch-site models in PAML (35). We found evidence of positive selection in 954 genes (LRT test FDR<0.1 in at least one branch-site test, Table S2) and many instances in which molecular convergence and positive selection were detected in the same gene (Fig. 3A; Fig. S3). In total, 403 genes showed at least one molecular convergence event between vertically transmitting lineages as well as positive selection in at least one of the lineages (Table S3).

**Figure 3.**
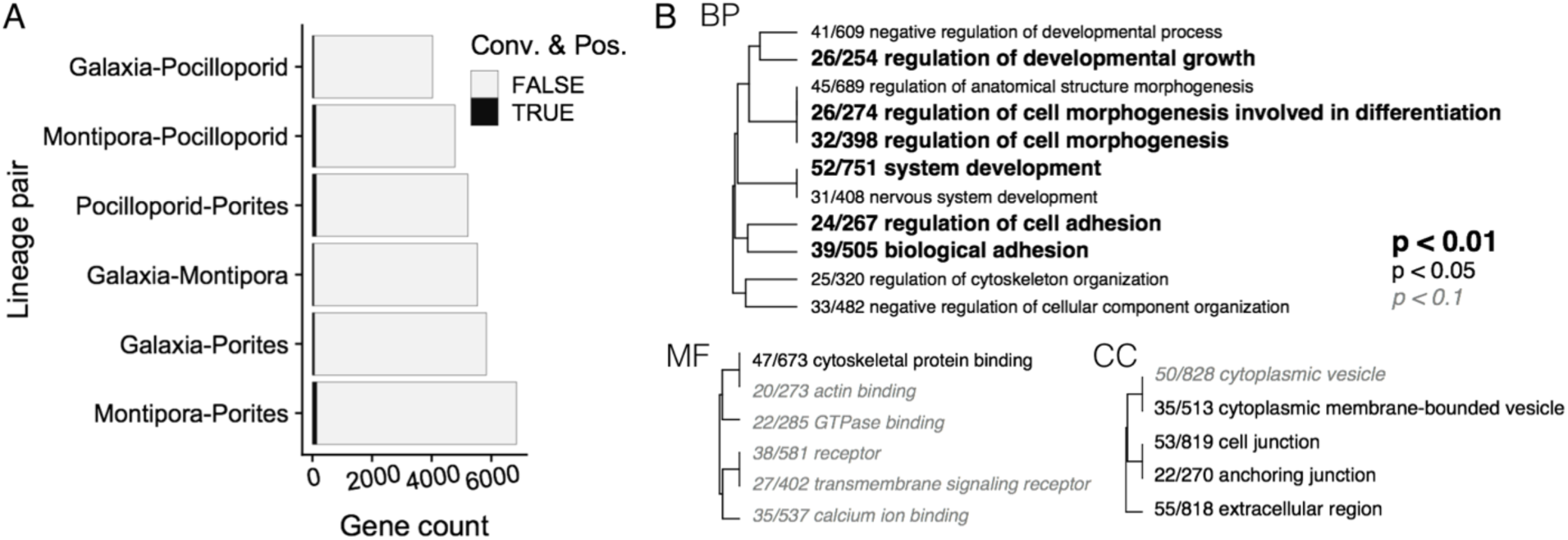
Frequency of genes exhibiting overlap in convergence and positive selection, and results of a categorical functional enrichment analysis of these candidates. (A) Frequency of genes exhibiting both signatures of convergence and positive selection per pair of vertically transmitting clades. Black shading indicates the set of genes with at least one convergence event and evidence of positive selection (FDR < 0.1) in at least one of the indicated lineages. (B) Gene ontology enrichment across all convergent and positively selected genes identified for any pair of vertically transmitting clades relative to the global gene list. Significance level is indicated by bolded text. (BP) Biological Processes, (CC) Cellular Component, (MF) Molecular Function.

Finally, we took advantage of the fact that the branch site test identifies individual amino acid positions that show evidence of positive selection (37), and identified a list of 100 genes for which the particular convergence event also showed evidence of positive selection in one or both lineages (branch site LRT p-value < 0.05 and BEB > 0.8; Table S4). No ontology enrichments were detected for this reduced group, but annotations were recovered for 66 of the 100 genes.

### The frequency of molecular convergence

The probability of parallel molecular evolution in response to selection is predicted to be twice as high as that under neutrality (38). Enforcement of vertical transmission in a laboratory manipulation of an anemone-*Symbiodiniaceae* symbiosis resulted in a host growth advantage, suggesting that the evolution of vertical transmission in Cnidarian symbioses may be favored by selection (39). However, an earlier analysis of genomic convergence among phenotypically convergent marine mammal lineages revealed that convergence was actually highest in terrestrial sister taxa in which no phenotypic convergence was evident, suggesting that the options for adaptive evolution may be limited by pleiotropic constraints (22). To assess the relative frequency of molecular convergence in our dataset we compared the proportion of molecular convergence in overlapping substitutions among three sets of phenotype pairs (vertical transmitters with other vertical transmitters, verticals with horizontals, and horizontals with other horizontals). This helped to control for possible confounding factors such as differences in mutation rate, and varying representation for each species based on data quality that may influence the absolute levels of molecular convergence detected (40).

We found no significant differences among phenotypic pairings in the mean proportion of molecular convergence (Fig. S4), molecular convergence and positive selection (Fig. S3), or specific convergence events in which the sites were also identified as being positively selected (Fig. S5). However, the proportion of convergence events is qualitatively different, and for each of these three data subsets, is higher among vertically transmitting pairs (Figure S3-S5). Although this pattern is tenuous, likely attributable to the small number of possible vertical-vertical comparisons, it is consistent with a proposed role of natural selection in driving the evolution of these convergent transmission mode phenotypes (5).

### Functional enrichments among top candidate genes

Coral symbionts reside within host gastrodermal cells, surrounded by a host-derived membrane known as the symbiosome (41). Although the specific genes mediating the establishment and long-term maintenance of this relationship remain unresolved, a number of biological processes are thought to be involved including host-microbe signaling, regulation of the host innate immune response and cell cycle, phagocytosis, and cytoskeletal rearrangement (6). To evaluate whether any of these previously highlighted processes were enriched among the 403 genes exhibiting both signatures of selection and convergent evolution, gene annotations were obtained from comparisons against the UniProt Swiss-Prot database (27) and a categorical functional enrichment analysis (FDR<0.1) was performed. Top functional enrichments (FDR<0.01) among biological processes (BP) terms included regulation of developmental growth and cell morphogenesis, and biological adhesion (GO:0048638; GO:0010769; GO007155; GO0022610). Endocytosis (GO:0006897) and immune response (GO:0006955) were also significant (FDR<0.1). Among molecular functions (MF), cytoskeletal protein binding (GO:0008092) was the most significant enrichment (FDR=0.016, Fig. 3B). Extracellular region (GO:0005576) was the most significantly enriched term among cellular components (CC), however, this term was also highlighted in a comparison of horizontally transmitting sister clades (Fig. S6), suggesting that it may be under selection in all corals and not necessarily specific to the evolution of vertical transmission. Additional top CC enrichments (FDR<0.1) specific to vertically transmitting lineages include cell junctions (GO:0030054) and cytoplasmic membrane-bounded vesicles (GO:0016023).

Three individual genes, ABL proto-oncogene 1 (ABL 1, ORTHOMCL8234), filamin C (ORTHOMCL8658), and poly(rC) binding protein 2 (ORTHOMCL8545), warrant additional discussion as they are classified among significantly enriched GO terms in all three ontology categories (BP, CC and MF) and were also among the less than 1% of genes in which the particular convergence event also showed evidence of positive selection (Fig. 4; Table S4). Importantly, none of these candidates have been previously implicated in the host-symbiont relationship in earlier analyses focusing on either coral bleaching, the breakdown of the symbiosis (42–46), or on the establishment of symbiosis in horizontally-transmitting corals (47– 49), highlighting the value of the present approach for identifying novel candidate genes potentially underpinning the coral symbiosis.

**Figure 4.**
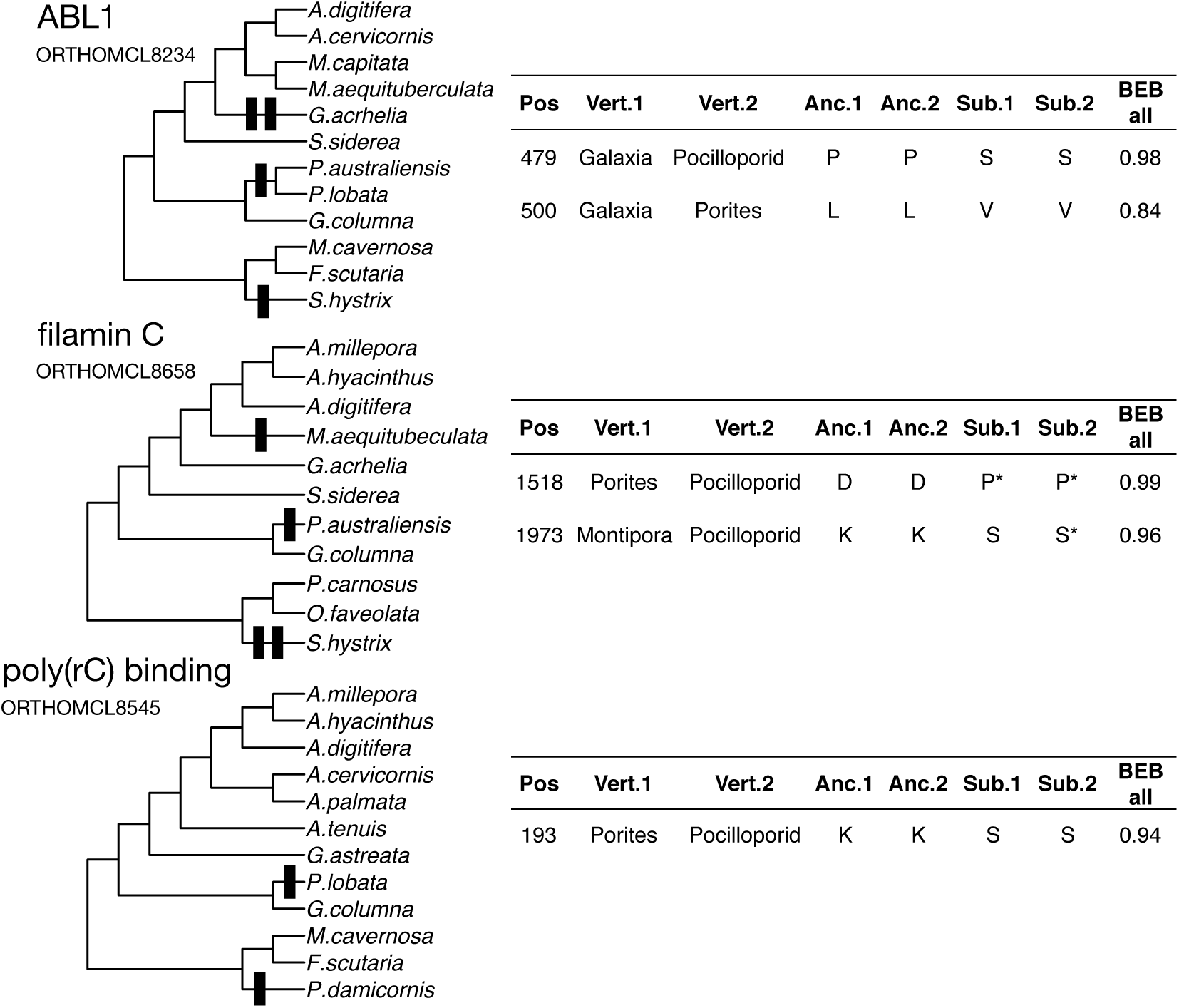
Select genes showing molecular convergence and positive selection at the same site. Left panels show gene trees constructed from nucleotides for each gene. Molecular convergence events that also showed evidence of positive selection are indicated with vertical bars. Tables show details of the molecular convergence events and evidence of positive selection: (Pos) amino acid position of convergence event; (Vert.1) first vertical lineage; (Vert. 2) second vertical lineage; (Anc.1) Ancestral amino acid for first vertical lineage; (Anc.2) Ancestral amino acid for second vertical lineage; (Sub.1) derived amino acid for first vertical lineage; (Sub.2) derived amino acid for second vertical lineage; (BEB all) Bayes Empirical Bayes posterior probability for positive selection at the position for the branch site test including all vertical transmitting lineages as foreground. Derived amino acids with BEB posteriors > 0.8 for tests using individual lineages as foreground are indicated with asterisks.

ABL 1 is a ubiquitously expressed nonreceptor tyrosine kinase known to be involved in organismal responses to a multitude of signals, including cell adhesion, DNA damage, oxidative stress and cytokines (50). This gene that has likely evolved to serve a variety of context-dependent biological functions, but is known to regulate several immune response phenotypes in mammals including antigen receptor signaling in lymphocytes, and bacterial adhesion to host cells (51–53). Through its role in regulating actin polymerization, ABL 1 is also involved in endocytosis (54), supporting the hypothesis that it may play a key role in mediating the heritable transmission of symbionts. Filamins are another family of actin-binding proteins which also exhibit great functional diversity in their interactions (55). While Filamin C was not identified in earlier functional genomic studies, expression of Filamin A was recently reported to be modified by temperature over the course of a monthly reproductive cycle in *Pocillopora damicornis*, a vertically-transmitting brooding coral (56). Similarly, Filamin B was found to be differentially expressed between symbiotic and aposymbiotic *Aiptasia* anemones (57). Combined, these results suggest an important role for this gene family in the maintenance and transmission of symbionts. Poly(C)-binding proteins also exhibit substantial functional diversity, but they are involved in transcriptional and translational regulation in addition to acting as structural components in DNA-protein complexes (58). Interestingly, poly(rC) binding protein 2 is a negative regulator of mitochondrial antiviral signaling protein (MAVS), a critical component of innate antiviral immunity, where overexpression has been shown to reduce, and knockdown to increase, cellular responses to viral infection (59). MAVS interacts with RIG-I-like (RLR) pattern recognition receptors, which are located in the cytoplasm, to identify foreign RNA (60). However, they have also been shown to function in defense against some bacterial pathogens (60, 61), suggesting that regulation of poly(rC) binding protein 2 could be involved in suppressing host innate immune responses against intracellular symbionts.

## Conclusions

Climate change and other anthropogenic processes threaten corals because of the sensitivity of the coral-dinoflagellate symbiosis to environmental stress (62, 63). Significant work has gone into investigating the breakdown of this relationship in the process known as ‘coral bleaching’ over the past three decades, yet fundamental questions remain unresolved, including a complete understanding of the genomic architecture underpinning the host-symbiont relationship (6, 64). Here, rather than asking about molecular mechanisms correlated with the breakdown of the coral symbiosis, we investigated a factor predicted to reinforce it: the evolution of vertical symbiont transmission. While the genes identified here represent promising candidates for further study, it is important to note that they likely represent only a fraction of the molecular changes involved in the evolution of symbiont transmission mode as there are alternate pathways to achieve the same phenotypic outcome that do not require changes at the level of the coding sequence (65). Increasing genomic resources will facilitate a deeper understanding of such alternative mechanisms, and the concurrent development of more advanced genetic tools for manipulating the coral (66) and other Cnidarian model symbioses (67, 68) will facilitate quantification of the precise phenotypic effects of these novel genes, as well as of changes in their sequence, contributing to a greater understanding of the cellular and molecular mechanisms underpinning this specific relationship, and necessary for the evolution of a heritable symbiosis.

## Methods

### *Sample preparation and sequencing for* Montipora aequituberculata *reference transcriptome*

Samples of *Montipora aequituberculata* were collected under the Great Barrier Reef Marine Park Authority permit G12/35236.1 and G14/37318.1. To generate a *M. aequituberculata* reference transcriptome, five replicate fragments of a single coral colony were subject to a two-week temperature stress experiment as described in (5) and snap frozen samples from control (27°C, days 4 and 17) and heat (31°C, days 2, 4 and 17) treatments were crushed in liquid nitrogen and total RNA was extracted using an Aurum Total RNA mini kit (Bio-Rad, CA). RNA quality and quantity were assessed using the NanoDrop ND-200 UV-Vis Spectrophotometer (Thermo Scientific, MA) and gel electrophoresis. RNA samples from replicate fragments were pooled in equal proportions and 1.8 µg was shipped on dry ice to the Genome Sequencing and Analysis Facility (GSAF) at the University of Texas at Austin where Illumina TruSeq Stranded libraries were prepared and sequenced on one lane of the Illumina Hiseq 4000 to generate 2 × 150 PE reads.

### Transcriptome assembly and annotation

Sequencing yielded 98 million raw PE reads. The *fastx_toolkit* (http://hannonlab.cshl.edu/fastx_toolkit) was used to discard reads < 50 bp or having a homopolymer run of ‘A’ ≥ 9 bases, retain reads with a PHRED quality of at least 20 over 80% of the read and to trim TruSeq sequencing adaptors. PCR duplicates were then removed using a custom perl script (https://github.com/z0on/annotatingTranscriptomes). Remaining high quality filtered reads (37.7 million paired reads; 6.7 million unpaired reads) were assembled using Trinity v 2.0.6 (69) using the default parameters and an *in silico* read normalization step at the Texas Advanced Computing Center (TACC) at the University of Texas at Austin. Since corals are ‘holobionts’ comprised of host, *Symbiodiniaceae* and other microbial components, resulting assemblies were filtered to identify the host component following the protocol described in (70).

### Additional transcriptomic resources

Transcriptomic data from 25 species of Scleractinia (stony corals) and 3 species of Actiniaria (anemones) were downloaded from the web (Table S1; (71); (72); (73); (74); (75); (42); (76); (77); (78); (79); (80); (81); (82); (83); (84); (70)).

### Protein sequence prediction

To prepare sequences for protein sequence prediction, we first modified sequence definition lines for each transcriptome to include the species name and an arbitrary sequence number. To remove highly similar isoforms, we used cd-hit (85) to cluster sequences with a sequence identity threshold of 0.98, alignment coverage for the longer sequence at least 0.3 and alignment coverage of the shorter sequence at least 0.3. For each resulting cluster, we retained only the longest sequence.

Protein coding sequences were predicted from the transcriptomic data based on open reading frames and sequence homology to known proteins and protein domains. Protein prediction steps were implemented with Transdecoder (86). First, the longest open reading frames (ORFs) were identified using a minimum amino acid length of 100. Then protein sequences were predicted from the longest ORFs based on blastp alignments against the Swissprot database (27) and protein domains identified with scanHmm in HMMER version 3.1b2 (28). The resulting coding sequence predictions were used for all downstream analyses. The predicted protein and coding sequences are available on github: https://github.com/grovesdixon/transcriptomes_convergent_evo_coral.git.

### Ortholog assignment

Predicted coding sequences were assigned to orthologous groups using FastOrtho, an implementation of OrthoMCL (29) available through Pathosystems Resource Integration Center (PATRIC) web resources (87)(http://enews.patricbrc.org/fastortho/). We ran FastOrtho using reciprocal blastp results with an e-value cutoff of 1e-10, excluding hits with alignment lengths less than 75% of subject sequences.

### Construction of species tree

To construct a species tree, we used a subset of 1,196 single-copy orthologous groups with at least 20 of the 28 taxa represented. The codon sequence alignments were concatenated in phylip format for input into RAxML (88). The species tree was generated with the rapid bootstrapping algorithm (100 iterations) using the GTRGAMMA model and three anemone species were used as an outgroup. Trees were visualized using Dendroscope (89) and Figtree http://tree.bio.ed.ac.uk/software/figtree/.

### Paralog pruning

When putative paralogs from the same taxon were monophyletic, all but the longest sequences were removed. This was done for an initial set of 20,563 orthologous groups for which at least 7 (25%) of the species were represented. Protein sequences for these orthologs were aligned with MAFFT using localpair (33) and gene trees were constructed using FastTree (90). At this point, sequences from the three anemone species were removed, and were not used for any further analyses. We used the biopython module Phylo (91) to identify gene trees for which multiple sequences from single species formed monophyletic groups. Removal of these sequences allowed us to include many more orthologous groups as single-copy orthologs (9,794 single copy orthologs prior to pruning, 13,092 after pruning). After pruning, putative single-copy orthologs were reverse translated into codon sequences using Pal2Nal (34).

### Phylogenetic ortholog filtering

Orthologous groups were further quality filtered based on monophyly of known clades. Here we constructed gene trees from nucleotide alignments of each single-copy ortholog. We checked these trees for monophyly of known clades (Genus *Acropora*, Genus *Montipora*, Genus *Galaxia*, Genus *Porites*, favid clade with *F. scutaria* as outgroup, pocilloporid clade with *M. auretenra* as outgroup, complex corals, robust corals), which were corroborated in our species tree (Fig. 1). For 58% of gene trees, all species fell within their expected clades. If a single sequence fell outside of its expected clade or clades, that sequence was removed and the ortholog was retained (27% of orthologous groups). If more than one sequence fell outside its expected clade, the ortholog was removed (15% of orthologous groups).

### Ancestral reconstruction and identification of convergent substitutions

We used ancestral reconstructions to infer molecular convergence for the remaining high-quality orthologous groups. For each orthologous nucleotide alignment, the ancestral amino acid was identified at each node in the species tree, as well as the amino acid changes that occurred along the branches of the tree. This analysis was performed with PAML (35), using the species tree as a the guide. Example control files are available on the Github repository (https://github.com/grovesdixon/convergent_evo_coral).

From the ancestral reconstruction results, we identified all substitutions that occurred at the same positions in two or more selected lineages (overlapping substitutions). The selected lineages included the branches leading to the common ancestor of four vertical transmitting clades, and their corresponding horizontally transmitting sister clade (eight clades total, Fig. 1). The horizontally transmitting sister clades were included to serve as negative controls, and for normalization of GO enrichment analyses (see below). In cases where a clade was represented by a single species, the terminal branch was used as the lineage for that clade (e.g. the two *Galaxia* species, Fig. 1).

Following (36), we considered both parallel and convergent substitutions as molecular convergence. For a given amino acid position, parallel substitutions refer to independent changes to the same amino acid from the same ancestral amino acid. Convergent substitutions refer to independent changes to the same amino acid from different ancestral amino acids. We also recorded all other types of independent changes at the same site (i.e. changes to different amino acids from the same ancestral amino acid, and changes to different amino acids from different ancestral amino acids).

### Testing for evidence of positive selection

We tested for evidence of positive selection using the branch-site test in PAML (35). Branch-site tests were performed on each ortholog using codeml with NSsites set to 2 and fix omega set to 1 for the null model, and set to 0 for the alternative model. Example command files and tree files are available on Github (https://github.com/grovesdixon/convergent_evo_coral). When labeling branches tested for evidence of positive selection for a given clade, only the branch leading to the most recent common ancestor of the clade was labeled (Fig. S7). In other words, whenever a vertically transmitting clade had more than one species, we tested for evidence of positive selection in the lineage leading to the common ancestor of the clade, rather than the terminal branches leading to each individual species. We made this choice because it seems likely that mutations enabling a vertical transmission phenotype occurred in the lineage leading to the common ancestor of the clade, in which vertical transmission was presumed to have already evolved. As with the convergence analysis, in cases where a clade was represented by a single species, the terminal branch for that species was labeled as foreground. Branch-site tests were performed for each individual clade, and for all vertically transmitting clades at once. Significance was tested using likelihood ratio tests, and p-values were adjusted to control for false discovery rate using the Benjamini-Hochberg procedure (92). As with our analysis of molecular convergence, we repeated the tests for the horizontally transmitting sister clades to serve as a negative controls and normalization of GO enrichment. It should be noted that a significant result for the branch-site test does not prove that positive selection occurred, it merely provides evidence that it may have occurred. For simplicity, we refer to genes significant for these tests as “positively selected” as in (22).

### Annotation of genes of interest

Genes of interest were selected based on an overlap in both evidence of positive selection and convergent substitutions. Genes were annotated based on the SwissProt database and Pfam hits used for protein prediction (e-value < 1e-5, and default parameters for hmmscan). Gene Ontology (GO) associations were applied to each orthologous group based on all SwissProt genes used for prediction of any of its constituent sequences. The GO annotations for these genes were gathered from the Gene Ontology Annotation (GOA) Database (93) ftp://ftp.ebi.ac.uk/pub/databases/GO/goa/UNIPROT/). For cases when sequences in an orthologous group were predicted with multiple different SwissProt hits, the orthologous group was annotated with GO associations from all included SwissProt genes. Some orthologous groups had only Pfam hits. These did not receive GO annotations.

### GO enrichment

GO enrichment was performed using Fisher’s exact tests on the final set of genes exhibiting overlap in evidence of positive selection in at least one of the branch site tests and had at least one molecular convergence event among the vertically transmitting lineages. A paired control analysis was performed for genes with the same signatures among the horizontally transmitting lineages (Fig. S6). To perform fewer total tests, and reduce the effect of false discovery correction, only large GO terms, associated with at least 200 orthologs in our dataset, were tested for enrichment.

## ACKNOWLEDGEMENTS

This work was supported in part by an NSF International Postdoctoral Research Fellowship, DBI-1401165 to CDK. Bioinformatic analyses were carried out using computational resources of the Texas Advanced Computer Center (TACC).

## STATEMENT OF AUTHORSHIP

CDK designed research and assembled new reference transcriptome; GBD analyzed convergence and selection; CDK wrote the first draft of the manuscript and both authors contributed to revisions.

## SUPPLEMENTARY MATERIAL

**Table S1.**
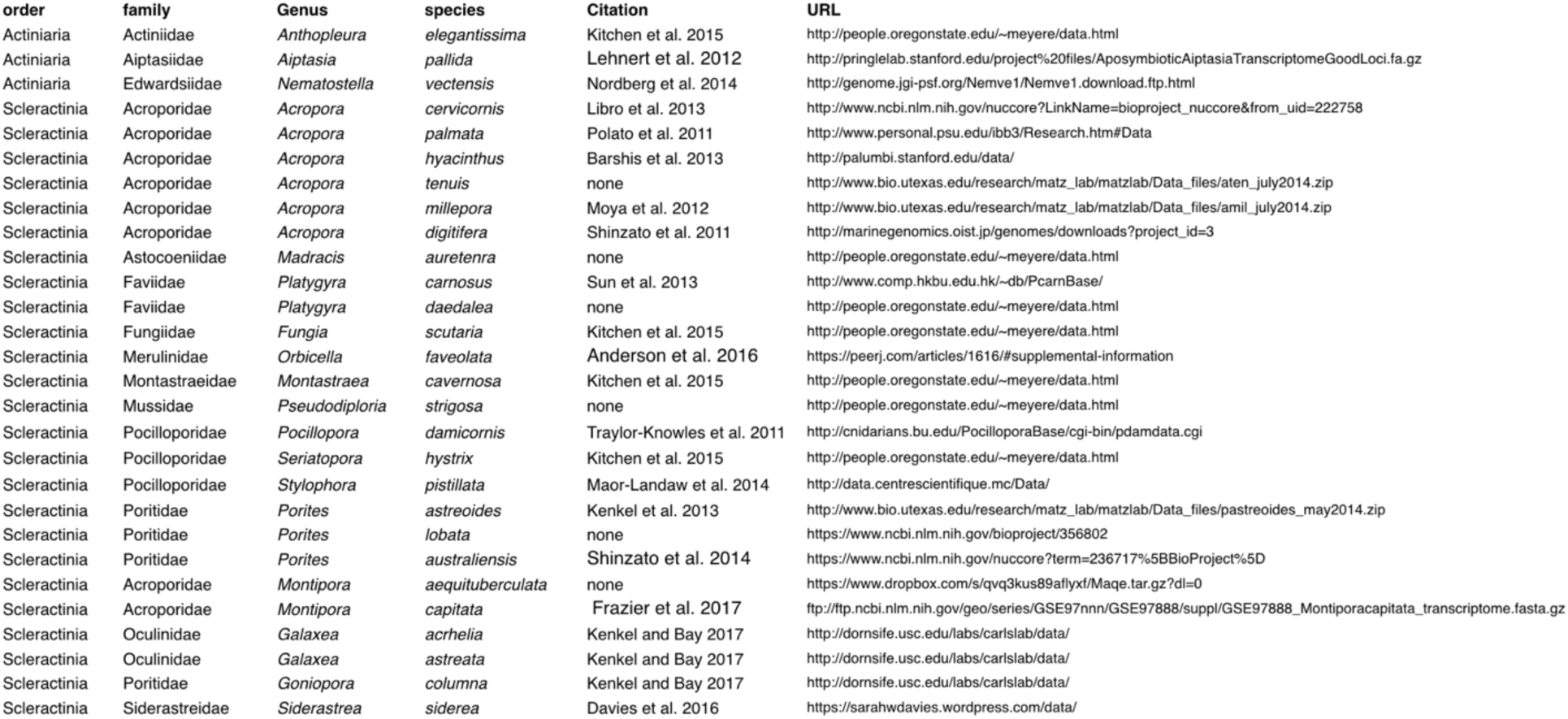
Sources of reference transcriptomes used for each species.

**Figure S1:**
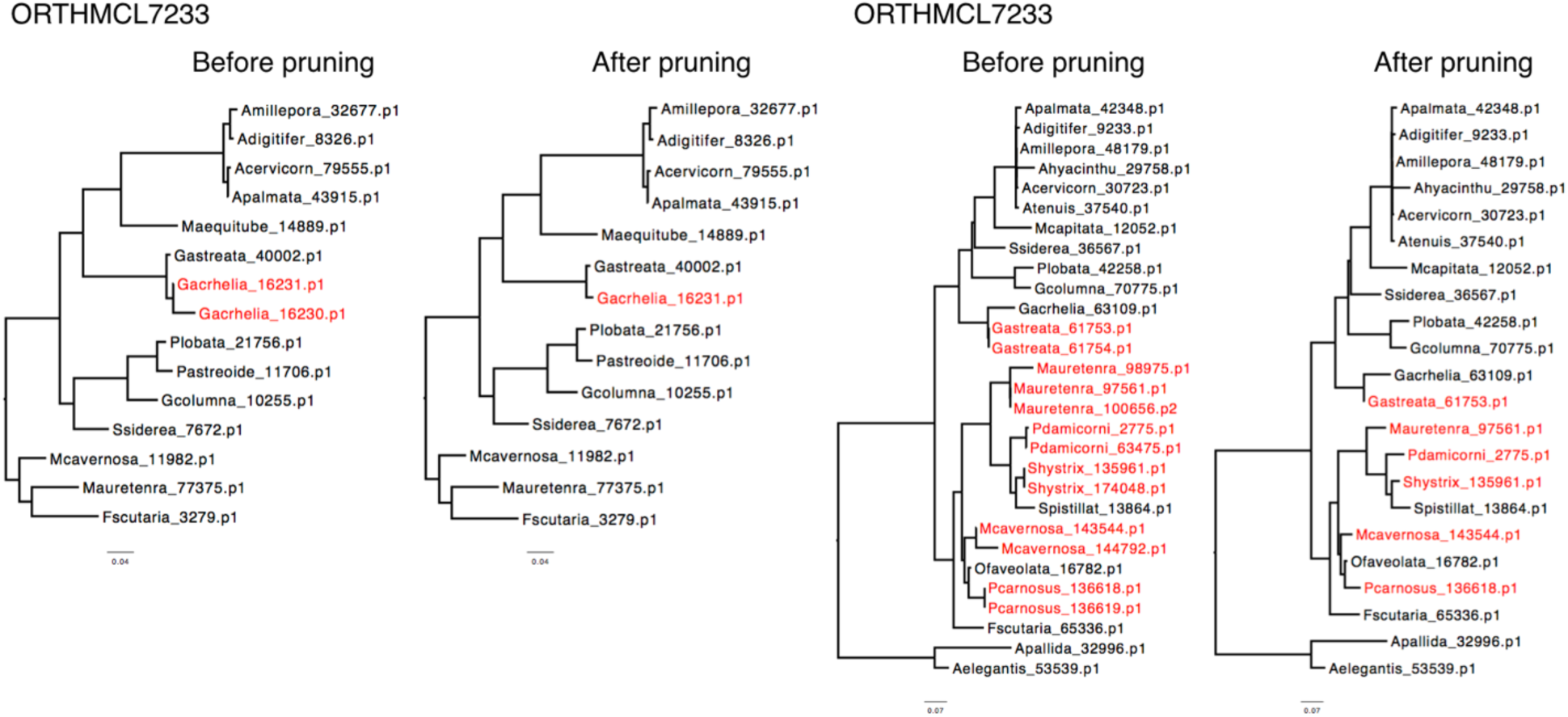
Examples of gene trees constructed for orthologous groups before and after paralog pruning. Paralog pruning was performed to remove duplicate sequences from orthologous groups if they came from a single species and formed a monophyletic clade. The figure shows gene trees for two different orthologous groups before and after pruning. Duplicated sequences from single species are shown in red. In the left orthologous group (ORTHMCL7233) a single duplicated sequence from *Galaxia acrhelia* was removed. The longer of the two sequences (Gacrhelia_16231.p1) was retained. In the right orthologous group, duplicate sequences formed monophyletic clades six species. In each of these cases, only the longest sequence was retained.

**Figure S2:**
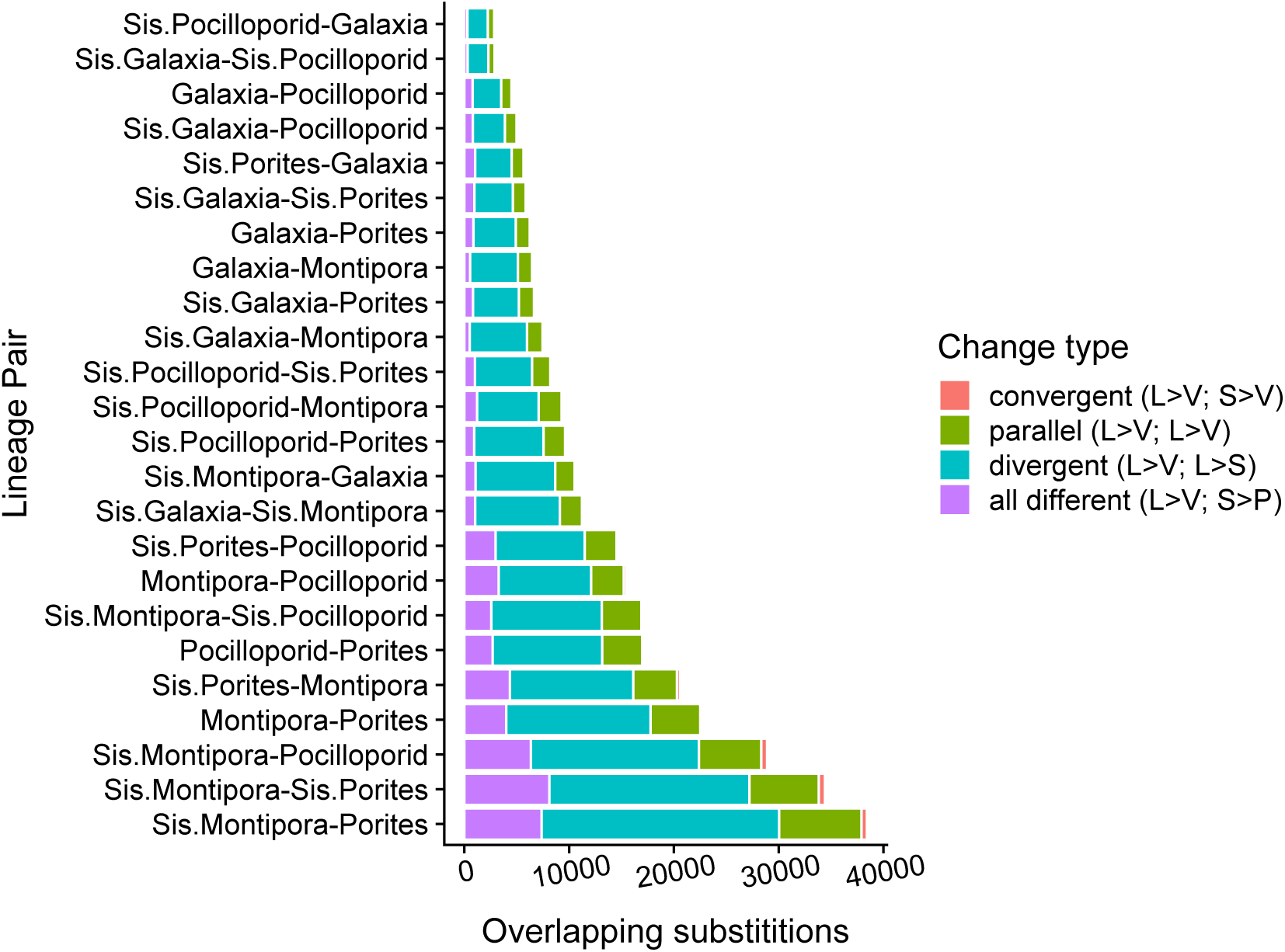
Categorization of all overlapping amino acid substitutions observed between all tested lineage pairs. An overlapping substitution is defined as an inferred amino acid change that occurred at the same position independently in the lineages leading to the common ancestor of the two indicated clades. To simplify comparisons, horizontal clades are labeled based on sisterhood to clades with vertical transmission (Fig. 1). Each overlapping substitution was classified into one of four categories: convergent substitutions (least frequent; salmon) are changes from different amino acids to the same amino acid; parallel substitutions (second most frequent; green) are changes from the same amino acid to the same new amino acid; divergent substitutions (most common; teal) are changes from the same amino acid to a different one; ‘all different’ substitutions (third most common; purple) are changes from different amino acids to different new amino acids. Examples of each type of overlapping substitution are show in in the legend.

**Figure S3:**
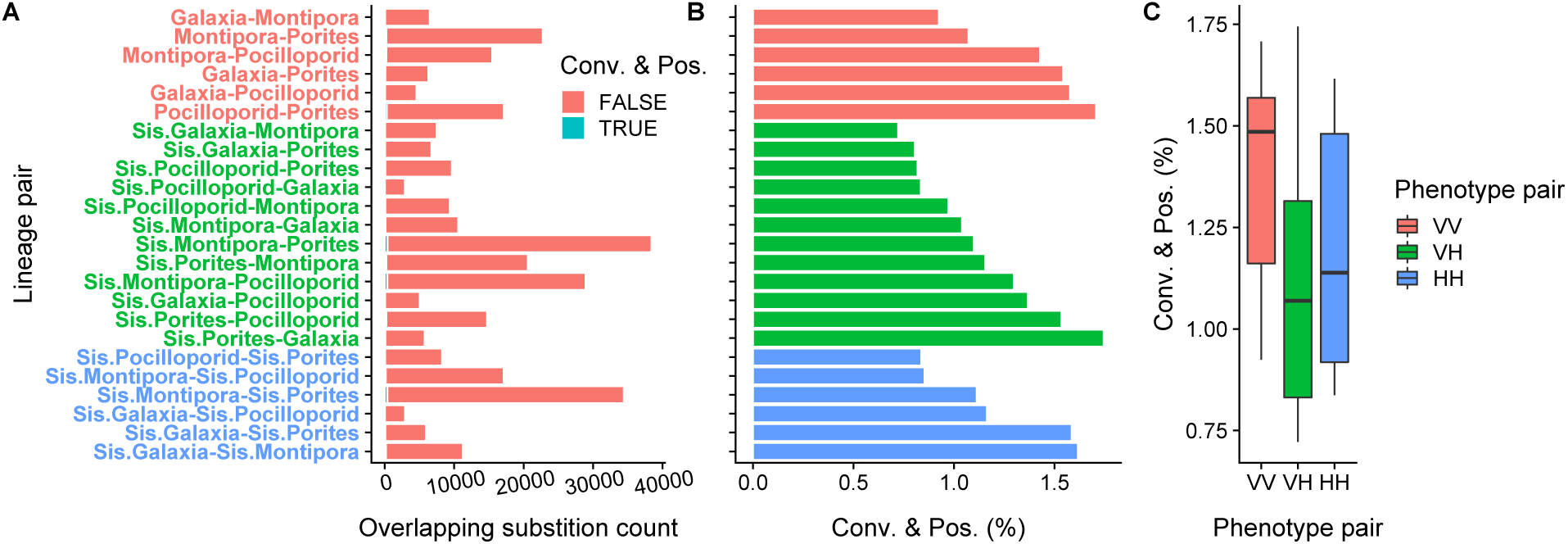
Comparison of frequency of convergent events among genes showing evidence of positive selection. (A) Counts of convergence events in genes showing evidence of positive selection in one or more of the indicated lineages. (B) Percentage of overlapping substitutions that were convergence events in genes also showing evidence of positive selection one or more of the indicated lineages. (C) Boxplot of the percentages in (B) split by phenotype pair, VV: vertical-vertical pairs, VH: vertical-horizontal pairs, HH: horizontal-horizontal pairs.

**Figure S4:**
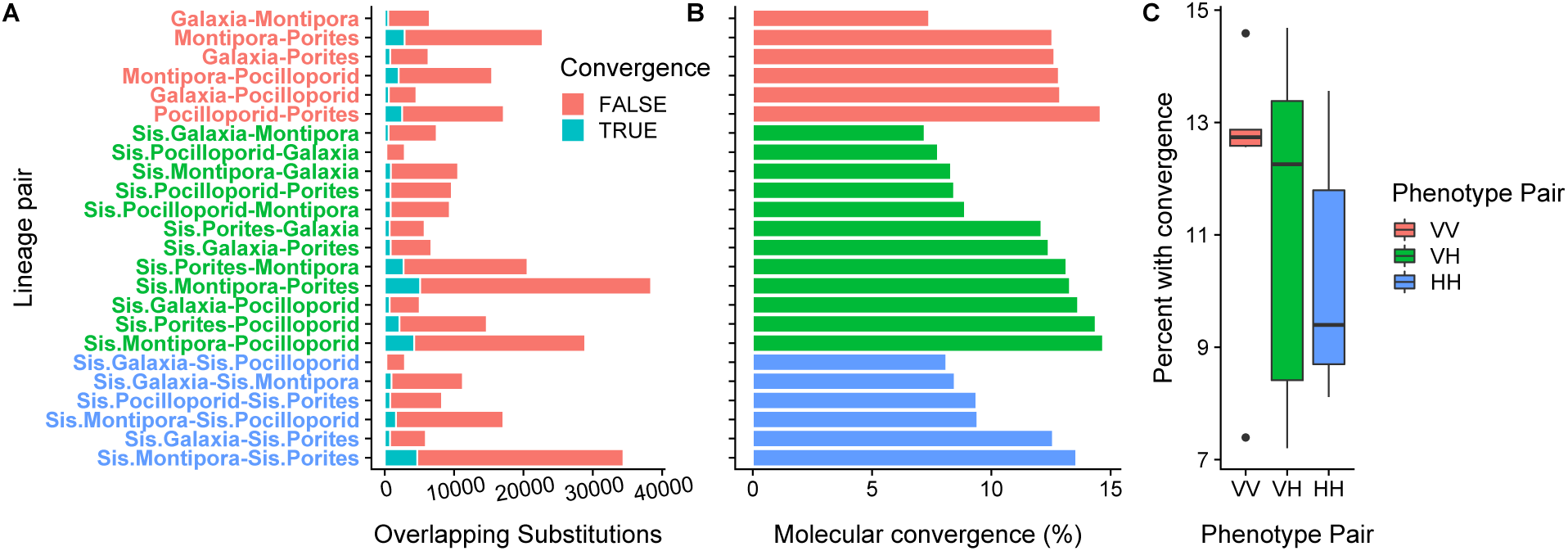
Comparison of the frequency of convergence events among overlapping substitutions. (A) Absolute counts of overlapping substitutions and convergence events for each species pair. (B) Percentage of overlapping substitutions that were convergence events. (C) Boxplot of the percentages in (B) split by phenotype pair, VV: vertical-vertical pairs, VH: vertical-horizontal pairs, HH: horizontal-horizontal pairs.

**Figure S5:**
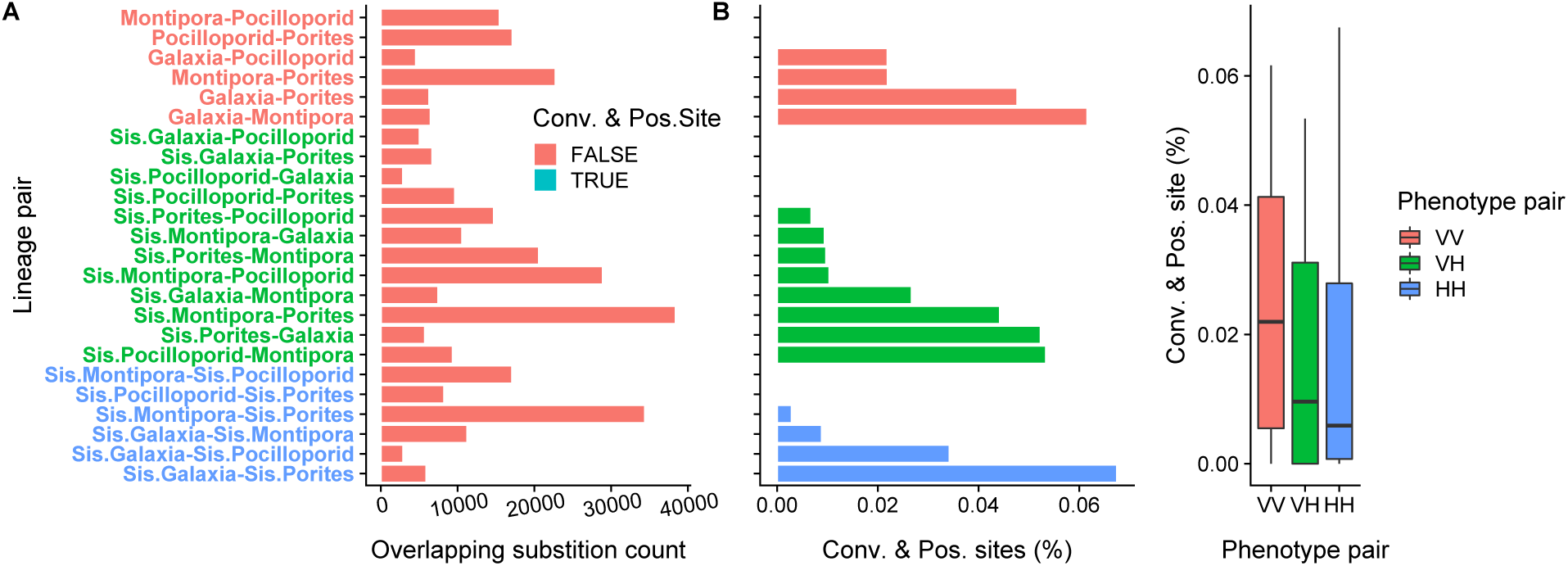
Comparison of the frequency of specific convergence events that were also identified as being positively selected. (A) Counts of convergence events which were also the sites exhibiting positive selection in one or more of the indicated lineages (Branch site test FDR < 0.1 for gene). (B) Percentage of overlapping substitutions that were convergence events in which the specific change was also the site of positive selection in one or more of the indicated lineages. Note that eight pairs have values of zero. (C) Boxplot of the percentages in (B) split by phenotype pair, VV: vertical-vertical pairs, VH: vertical-horizontal pairs, HH: horizontal-horizontal pairs.

**Figure S6:**
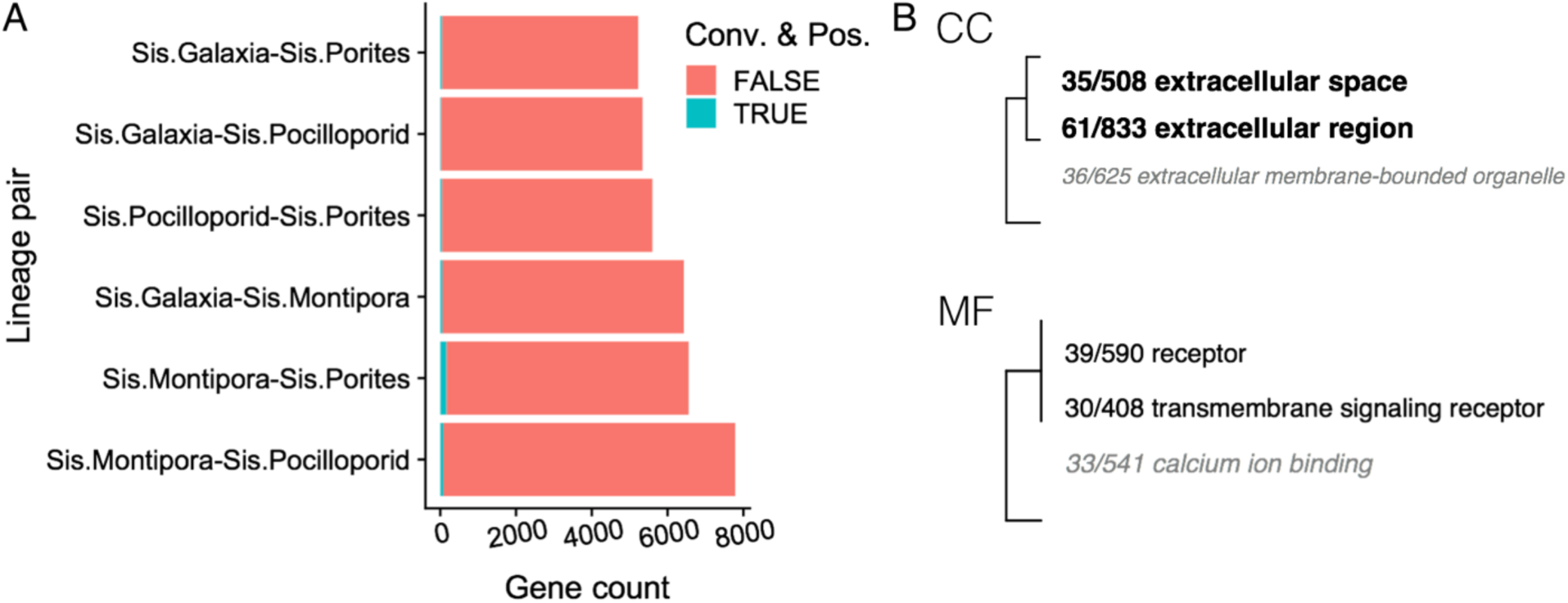
Functional enrichment for genes with convergence events and evidence of positive selection among horizontally transmitting sister clades. (A) Frequency of tested genes showing convergence and positive selection per pair of horizontally transmitting clades. Teal shading indicates the set of genes with at least one convergence event and evidence of positive selection (FDR < 0.1) in at least one of the indicated lineages. (B) Gene ontology enrichment across all convergent and positively selected genes identified for any pair of horizontally transmitting clades relative to the global gene list. Significance level is indicated by bolded text. Fractions preceding ontology terms indicate … (BP) Biological Processes, (CC) Cellular Component, (MF) Molecular Function. No ontology terms for Biological Process were significant.

**Figure S7:**
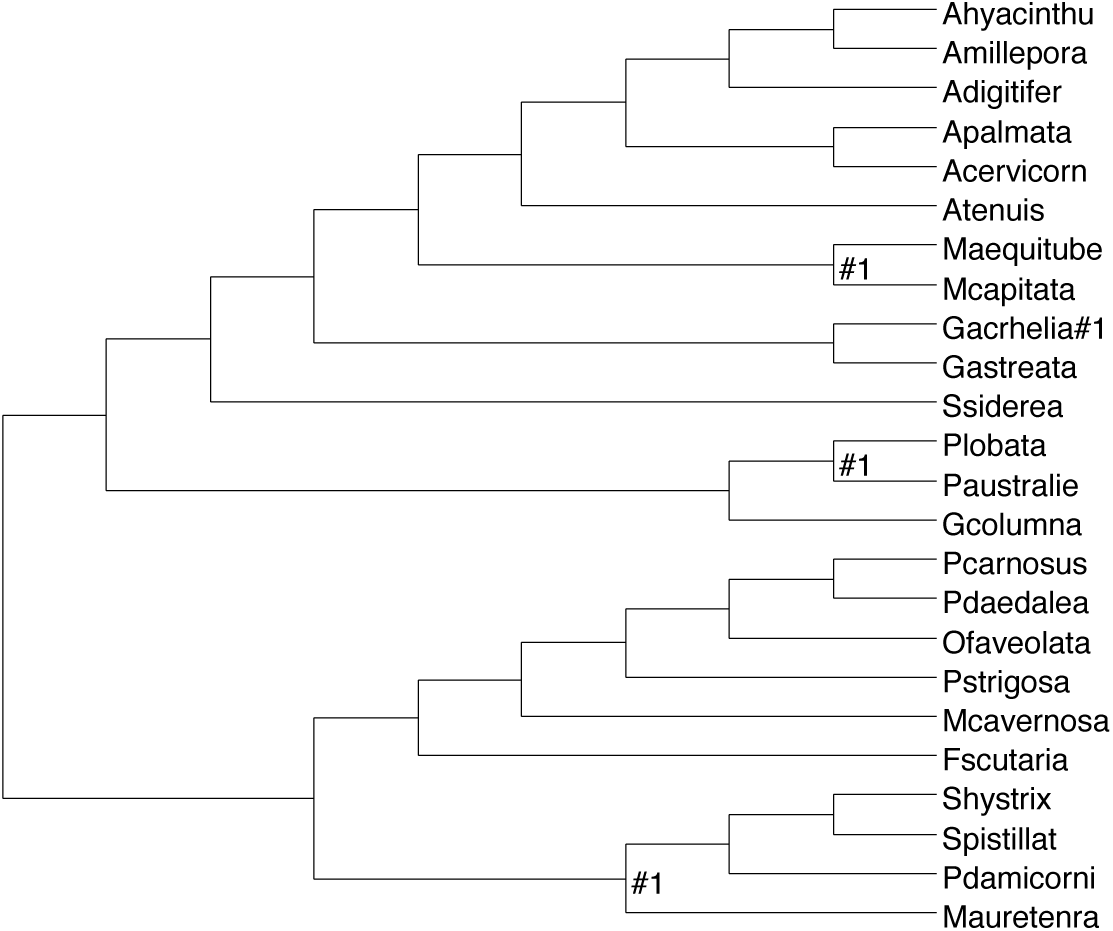
Labeling of branches for branch site tests. When performing the branch site test, the branch or branches being tested for evidence of positive selection are labeled with “#1”. When testing for evidence of positive selection in a clade, we labeled only the branch leading to the common ancestor of that clade. In cases when a clade had only a single species, for example *Galaxia acrhelia*, the branch for that species was labeled.

